# Interleukin-15 Receptor α Chain Knockout NOD-SCID mice have reduced numbers of NK cells, develop thymic lymphomas and fail to engraft human hematopoietic cells

**DOI:** 10.1101/2021.04.22.440980

**Authors:** Andrea Vecchione, Grace Nauman, Mohsen Khosravi Marharlooei, Nichole Danzl, Hao-Wei Li, Estefania Chavez, Benjamin E. Low, Michael V. Wiles, David Serreze, Lei Ding, Megan Sykes

## Abstract

NOD-SCID IL2rγ_c_^null^ (NSG) mice are widely used to model human immune cell development because they are more permissive for human hematopoietic cell engraftment and reconstitution than NOD-SCID mice. While increased human reconstitution in the blood of NSG mice has been attributed to the absence of mouse NK cells, deletion of the common gamma chain (γc) limits development of lymphoid tissue inducer cells and precludes development of normal secondary lymphoid structures. The disorganized lymphoid tissue leads to compromised human T cell-B cell interactions and results in variable human immune cell function in human immune system (HIS) NSG compared to NOD-SCID mice. We attempted to remove mouse NK cells from NOD-SCID mice while retaining other γ_c_-dependent cytokine responses by targeted disruption of the mouse genomic *IL15RA* locus with CRISPR/Cas9. IL15Rα is required for the development, function and survival of NK cells. NOD-SCID IL15Rα^-/-^ mice showed reductions in NK cells and NK cell function. However, NOD-SCID IL15Rα^-/-^ mice demonstrated accelerated thymic lymphomagenesis and showed earlier mortality compared to NOD-SCID mice. This result suggests that mouse NK cells are important to delay lymphoma development in NOD-SCID mice. We transplanted thymectomized NOD-SCID ILl5Rα^-/-^ mice with human fetal liver CD34^+^ cells and thymus to determine if these mice supported engraftment and development of a transplanted HIS. Surprisingly, we found that peripheral human engraftment was inferior (mean 0.05% of lymphocytes) to that in both NOD-SCID (mean, 10.5% of lymphocytes) and NSG (mean, 54% of lymphocytes) mice. These results indicate that NOD-SCID IL15Rα^-/-^ mice are not permissive for human CD34^+^ cell engraftment.

## Introduction

Immunodeficient mice transplanted with functional human cells and tissues, or “human immune system mice,” are a powerful in vivo model for studying human cells in a controllable, experimentally tractable context [1],[2]. HIS mice transplanted with human fetal liver CD34^+^ cells and fragments of human fetal thymus can repopulate with human CD45^+^ immune populations, including B cells, T cells and antigen-presenting cells (APCs) [1],[2]. Model development was initially hampered by rejection of transplanted human cells by the mouse immune system, including murine NK cells [3],[4]. Development of the NOD-SCID mouse strain, which lacks mouse T and B cells [5],[6] and expresses a murine SIRPα that interacts effectively with human CD47 to prevent murine macrophages from engulfing human cells [5]–[7], provided a significant advance. However, NOD-SCID mice develop thymic lymphoma, which limits their life span to around 30 weeks and makes the model unsuitable for long-term transplantation studies [8],[9]. Moreover, NOD-SCID mice have suboptimal human hematopoietic cell engraftment due to murine NK cell activity. Removal of the IL-2 receptor common gamma chain (*γ*_c_) to create the NOD-SCID IL2rγ_c_^null^ (NSG) strain, was a further step forward [10]–[12]. The *γ_c_* chain is required for high-affinity ligand binding and signaling through multiple cytokine receptors, including IL-2 and interleukins 4, 7, 9, 15, and 21 [13]–[16]. Disrupting signaling through these cytokine receptors leads to severe defects in both adaptive and innate immune cell development in the mouse [16]. Immunodeficient mice bearing a targeted mutation in the *IL2RG* gene support significantly higher levels of human hematopoietic and lymphoid cell engraftment than all previous immunodeficient strains of mice [17].

Despite these advances, HIS NSG mice show a limited ability to generate primary and recall humoral immune responses, which may be due to weak or inefficient human T cell-B cell interactions in lymphoid structures. This is evidenced by the observation that immunodeficient mice transplanted with human CD34^+^ cells do not mount consistent vaccination responses with class-switched antibodies [18]. This phenomenon may be partially explained by the lack of *γ*_c_ chain signaling, which leads to a decrease in production of mouse lymphoid tissue inducer cells and prevents robust development of secondary lymphoid structures [19]. With intact *γ*_c_ chain signaling, NOD-SCID mice develop secondary lymphoid structures such as cryptopatches in the gut [20], while NSG mice lack these structures. Cryptopatches in NOD-SCID mice served as anlagen for the development of human gut-associated lymphoid structures (GALT) in reconstituted NOD-SCID mice [1],[11],[20].

Because depletion of NK cells in NOD-SCID mice has been shown to successfully improve human hematopoietic cell engraftment [21], we aimed to develop a model that specifically lacks murine NK cells, but has otherwise intact common gamma chain cytokine receptors, by targeting the *IL15RA* locus with CRISPR/CAS9 technology. IL15Rα is an important component of the heterotrimeric plasma membrane receptor for the IL-15 cytokine [22]. IL-15 and IL-15Rα are required for NK cell development and function [23],[24]. They have also been shown to play important roles in skeletal muscle metabolic function [25] and bone formation [26]. We generated a knockout mouse strain that lacks IL15Rα on the NOD-SCID background, termed NOD-SCID IL15Rα^-/-^ mice. We found that NOD-SCID IL15Rα^-/-^ mice have fewer NK cells than their NOD-SCID littermates, but also observed more rapid development of thymic lymphoma and reduced lifespan compared to NOD-SCID mice. Since NK cells were reduced in these mice, we hypothesized that NOD-SCID IL15Rα^-/-^ mice might be more permissive hosts for transplanted human cells than NOD-SCID or NSG mice. However, when we transplanted thymectomized NOD-SCID IL15Rα^-/-^ mice with human fetal liver CD34^+^ cells and fetal thymus tissue from the same donor, we found that human engraftment was markedly reduced compared to that in NOD-SCID and NSG littermates. This unexpected observation suggests that murine IL15Rα, either directly or indirectly, plays a critical, previously undescribed role in supporting engraftment of transplanted human hematopoietic progenitor cells in immunodeficient mice. Since our studies show that murine NK cells delay lymphoma development in NOD-SCID mice, the NOD-SCID IL15Rα^-/-^ mouse could be used to help understand the role of NK cells in anti-tumor immunity.

## Materials and Methods

### CRISPR/CAS9

Guide RNAs 5’-GTCCCCAGCAGTGACCCAAC-3’ and 5’-GCCATGGCCTCGCCGCAGCTC-3’, targeting the start codon of the *IL15RA* locus (IL15Rα ATG), were designed using the sgRNA Designer, from Broad Genetic Perturbation Platform Web Portal (Cambridge, MA), available at https://portals.broadinstitute.org/gpp/public/analysis-tools/sgrna-design-help. Sequences were confirmed using the Wellcome Sanger NOD/ShiLtJ reference sequence (Hinxton, UK; https://www.sanger.ac.uk/science/data/mouse-genomes-project). gRNAs were purchased from Integrated DNA Technologies (Coralville, IA) and cloned into the pX335 plasmid (Addgene, plasmid #42335) encoding genetically modified Cas9 endonuclease which generates nicks, rather than double-strand breaks, in targeted DNA [27]. Using a nickase combined with multiple targeting gRNAs reduces off-target genomic mutations, a major concern with standard CRISPR approaches. gRNAs were validated in EL4, a murine T cell lymphoma^25^ that expresses IL15Rα. EL4 cells were propagated in RPMI 1640 (Thermo Fisher Scientific) supplemented with 10% heat-inactivated fetal bovine serum (FBS, Gemini BioProducts), 10mM HEPES (Gibco, Gaithersberg, MD), 20mM L-glutamine (ACROS Organics, Geel, Belgium), 1mM sodium pyruvate (Fisher BioReagents, Fair Lawn, NJ) and 0.2% 2-Mercaptoethanol (MP Biomedicals, LLC, Santa Ana, CA). gRNA_pX335 plasmids were transfected into EL4 cells by nucleofection using an Amaxa nucleofector and the Cell Line Nucleofector Kit L (Lonza, Basel, Switzerland) according to the manufacturer’s instructions. We verified genomic disruption of IL15Rα locus using the IDT Surveyor Mutation Detection Kit (Integrated DNA Technologies) per manufacturer’s instructions. We confirmed that gRNA targeting led to a reduction in surface IL15Rα expression by antibody staining 120hrs after transfection, using an anti-IL15Rα antibody (APC-labeled DNT15Rα, eBiosciences, San Diego, CA).

### Animal use

NOD-SCID IL15Rα ATG^4^ mice were generated at The Jackson Laboratory by Cas9/sgRNA microinjection in NOD-SCID embryos (stock #1303). Single guides (target sites: 5’-GTCCCCAGCAGTGACCCAAC and 5’-GCCATGGCCTCGCCGCAGCTC) were generated using in vitro transcription as described previously [28]. Cas9 mRNA (Trilink, 60ng/μl), Cas9 protein (PNABio, 30ng/μl), sgRNA (30ng/μl each), and RNAsin (Promega, 0.2U/μl), were prepared in RNAse-Free TE (10mM Tris/0.1mM EDTA) and delivered into fertilized NOD-SCID oocytes by pronuclear microinjection. Offspring were screened using PCR and Sanger sequencing. PCR Primers (FWD: 5’-TGCAGGGAAGACAGCATTAAG-3’, REV: 5’-AGAGGCACACTCACACATTTC), were designed to cover the IL15Rα Exon 1, generating a 785bp amplicon from the WT allele. We selected a line with 38bp of Exon 1 deleted (GTCCCGTTGGGTCACTGCTGGGGACAATTGGCC**<u>ATG</u>**GC), including the start codon. This line was assigned the JAX stock # 30327.

NOD-SCID IL15Rα full-/- animals were generated at The Jackson Laboratory by Cas9/sgRNA microinjection in NOD-SCID embryos (stock #1303) using sgRNAs targeting Exon 1 (5’-GCCGCAGCTCCGGGGCTA) and Exon 6 (5’-GCCTGGTACATCAAATCA), generated as described above. In an effort to facilitate the deletion of the intervening sequence, a 200nt oligonucleotide designed to bridge the cut sites was included, shown here:

(5’-AAGGGACCCCTCTATCGGGAGGGCCTCTTGGAATCGTTATGATCAAGCTTTTTTGGTAGAAAGGAAGCACTATGCCCTGCAGTTCTTCTGCTCACCTTGACCCCGGAGCTGCGGCGAGGCCATGGCCAATTGTCCCCAGCAGTGACCCAACGGGACGCAAGGCTGGGGTCCCTGGGAGGTGGCGAGCTCAGCTCCAGGAC). Cas9 mRNA (Trilink, 60ng/μl), Cas9 protein (PNABio, 30ng/μl), sgRNA (30ng/μl each), bridging oligonucleotide (3ng/μl) and RNAsin (Promega, 0.2U/μl), were prepared in RNAse-Free TE (10mM Tris/0.1mM EDTA) and delivered into fertilized NOD-SCID oocytes by pronuclear microinjection. Offspring were screened using PCR and Sanger sequencing. PCR Primers (FWD: 5’-TGCAGGGAAGACAGCATTAAG, REV: 5’-CAGAATCAAGGCTGACATGAAC) designed to cover the IL15Rα Exons 1 through 6, were predicted to generate a ~700bp amplicon from the mutant allele, while no product was expected from the WT allele (25,473bp). Ultimately, a line with 24,778bp deletion from Exon 1 through Exon 6 was established and characterized, the PCR product for this allele is 694bp. For reference, the flanking sequence (15bp on either side of the deletion) is shown here (5’-cgccgcagctccggg| tgagcagaagaactg), this line was assigned the JAX stock # 30326 (Supplementary Figure 1B-C-D).

Mice targeted at the ATG and the Exon 6 site were found to have a larger portion of the *IL15RA* locus disrupted, but NK cell phenotypes were indistinguishable.

Genetically modified mice were propagated at Columbia University Irving Medical Center in a *Helicobacter, Pasteurella pneumotropica* and Specific Pathogen-Free animal facility. In some cases, heterozygous knockout animals were provided and were crossed to NOD-SCID, some of whose progeny served as IL15Rα WT control animals. NSG animals were obtained from The Jackson Laboratory. All animal procedures were approved by the Institutional Animal Care and Use Committees of the Columbia University Irving Medical Center and The Jackson Laboratory.

### Human tissue transplantation

Human fetal thymus and fetal liver (FL) tissues (gestational age, 17 to 20 weeks) were obtained from Advanced Biosciences Resources. Fetal thymus fragments were cryopreserved in 10% dimethyl sulfoxide (Sigma-Aldrich) and 90% human AB serum (Gemini Bio Products). FL fragments were treated for 20 minutes at 37°C with 100 μg/mL of Liberase (Roche) to obtain a cell suspension. Human CD34^+^ cells were isolated from FL by density gradient centrifugation (Histopaque-1077, Sigma) followed by positive immunomagnetic selection using anti-human CD34 microbeads (Miltenyi Biotec) according to the manufacturer’s instructions. Cells were then cryopreserved as described [29]. Four-to eight-week old mice were thymectomized [30] and allowed to recover for two weeks, then sublethally irradiated (1.0 Gy) using an RS-2000 X ray irradiator (Rad Source Technologies, Inc., Suwanee, GA). Human fetal thymus fragments of about 1-3 mm^3^ were implanted beneath the kidney capsule and 2 × 10^5^ human FL CD34^+^ cells were injected intravenously. Recipient mice were treated IV with 400 μg of anti-human CD2 monoclonal antibody [29] (Bio X Cell, Inc) by intravenous injection on days 0, 7, and 14.

### Isolation of intrahepatic mononuclear cells

Livers were procured from humanely euthanized donor mice. After removal of the gall bladder, the liver was minced to 1 cm^3^ pieces. and dissociated to a single cell in RPMI-1640 (Thermo Fisher Scientific, Waltham, MA) media by gently pushing through a 100 μm steel sieve (Bellco). The non-homogenized tissue was pelleted by centrifuging the filtered liver samples for 1 minute at 50 x g. The single cells suspension in the supernatant was centrifuged for 5 minutes at 400 x g and then resuspended in 10 mL of 37.5% isotonic Percoll [29]. Density gradient centrifugation for 30 minutes at 850 x g separated a top layer containing cellular debris and a pellet containing mononuclear cells. The mononuclear cells were washed with RPMI-1640. The liver mononuclear cells were enriched with a second density gradient centrifugation over Histopaque-1077 according to manufacturer’s instructions (Sigma). The interface layer containing mononuclear cells was collected, washed and counted using Trypan Blue dye to exclude dead cells.

### Flow Cytometry (FCM)

Human reconstitution, mouse NK cells, and presence of murine thymic lymphoma cells were assayed in the peripheral blood of transplanted mice every two weeks. At the time of sacrifice, single-cell suspensions were prepared from the bone marrow (BM), spleen, liver, lymph nodes (LN) and blood for staining. Single-cell suspensions of BM, spleen and blood were treated with ACK lysis buffer (Life Technologies) to remove erythrocytes. Single cell suspensions of mouse livers were filtered through a 100μm cell strainer (BD Falcon, Franklin Lakes, NJ). Dead cells, hepatocytes and debris were removed from liver samples as described above. After ACK erythrocyte lysis, remaining cells were passed through a 40μm filter prior to FCM analysis. All cells were stained using the following mAbs; anti-hCD3 FITC (OKT3), anti-Ter119 PE-Cy7 (Ter119), anti-mCD45 APC-Cy7 (30-F11), anti-hCD45 PE-CF594 (HI30), anti-hCD19 PERCP-Cy5.5 (HIB19), anti-hCD4 V500 (RPA-T4), anti-hCD8 AF700 (RPA-T8), anti-hCD14 BV650 (M5E2), anti-mCD3 PE (SP34-2), anti-mCD4 PB (GK1.5), anti-mCD8 APC (53-6.7), anti-mB220 FITC (RA3-6B2), anti-mCD335 PERCP-efluor710 (29A1.4), anti-mDX5 biotin (DX5), anti-mThy1.2 biotin (53-2.1). Data were collected using a BD Fortessa cytometer and analyzed using FlowJo (TreeStar, Ashland, OR).

### NK cell cytotoxicity assay

To determine NK cell cytotoxicity, we adapted a previously described radioisotope-free method [31]. Briefly, RL♂1 murine class I-deficient lymphoma cells were cultured in RPMI 1640 supplemented with 10% fetal bovine serum (FBS, Gemini BioProducts, West Sacramento, CA) and used as target cells. Erythrocyte-depleted splenocytes from NSG mice were used as control cells. Control and target cell populations were labeled with fluorescent amine binding dyes CFSE (Molecular Probes, Eugene, OR) and e450 (eBiosciences) respectively, and combined in a 1 to 1 ratio. Since NK cells are the only cytotoxic populations present, bulk erythrocyte-depleted splenocytes from age and sex-matched NOD-SCID and NOD-SCID IL15Rα^-/-^ mice were harvested as effectors and mixed with the Target/Control mixture at the indicated ratios. Cells were incubated together in RPMI 1640 (Thermo Fisher Scientific, Waltham, MA) supplemented with 10% FBS for 4 hours at 37°C with 5% CO_2_. Then, cells were stained for viability with DAPI (Sigma Aldrich). Percentage of specific killing was calculated using the following formula:

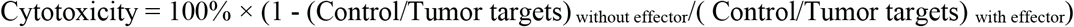

### Statistical analysis

Statistical analysis was performed with Graph-Pad Prism 6.0 (GraphPad Software). All data are expressed as average ± standard error of mean. The nonparametric Mann-Whitney U test was used to compare groups at individual time points for each cell population. Two-way analysis of variance (ANOVA) with post hoc Bonferroni test was used to compare 2 or 3 groups. The parametric paired t-test was used in the killing assay. Survival curves were compared using the Log-rank (Mantel-Cox) test. P < 0.05 was considered statistically significant.

### Results 1.1 Generation of NOD-SCID IL15Rα^-/-^ mouse model

We generated NOD-SCID IL15Rα^-/-^ mice using two sgRNAs to target the start codon (ATG) of the *IL15RA* locus (Figure 1A and Supplementary Figure 1A). Following CRISPR-mediated non-homologous end-joining (NHEJ) in the microinjected NOD-SCID embryos, a variety of alleles were generated. Of 19 mice born, 9 had clearly identified mutations at the target site, as evidenced by PCR and Sanger sequencing (data not shown). After back-crossing selected founders to NOD-SCID, the precise alleles in the N1 heterozygotes were characterized. Ultimately, a single line carrying a 38bp depletion was selected and bred to homozygosity (Supplementary Figure 1A). We validated the efficacy of these two sgRNAs by transfecting murine EL4 cells with plasmids containing gRNAs and performed Surveyor nuclease analysis on a PCR-amplified region targeted by the sgRNA to confirm that IL15RA start codon was disrupted (data not shown). In addition, IL15Rα surface expression was absent on the surface of EL4 cells (data not shown). Deletion of 38bp at the ATG site was detectable by PCR (Figure 1B). Similar to NSG mice, NOD-SCID IL15Rα^-/-^ mice demonstrated reduced numbers of NK cells in the blood, liver and spleen compared to NOD-SCID mice (Figure 1C and 1D).

**Figure 1.**
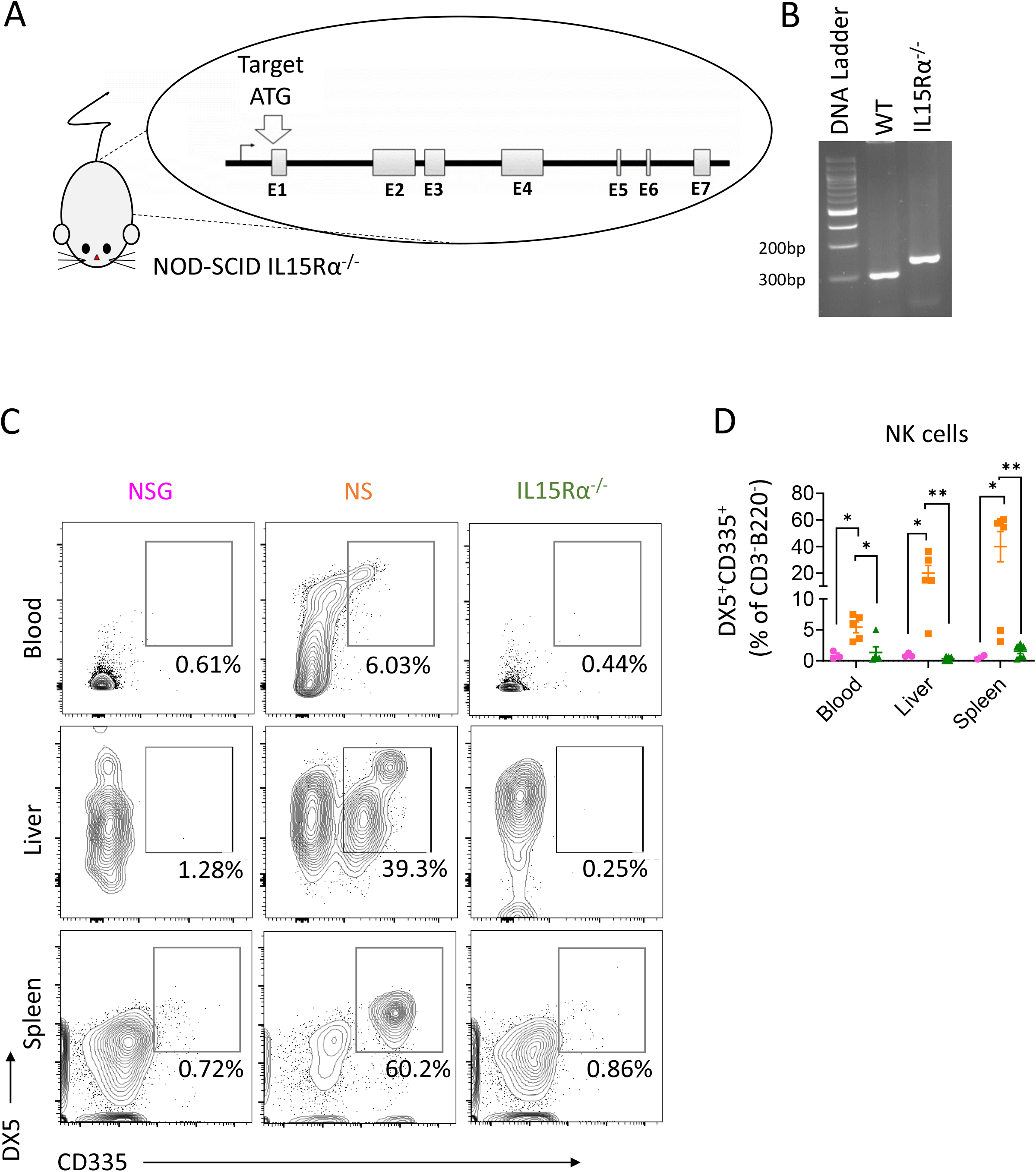
CRISPR-mediated targeting of the *IL15RA* locus reduces NK cells on a NOD-SCID background. **A.** Diagram of the *IL15RA* locus indicating the CRISPR target sites near the start codon (ATG). **B.** CRISPR-mediated 38bp deletion at the ATG of the IL15RA locus is detectable by PCR amplification of genomic DNA from NOD-SCID (WT) and NOD-SCID IL15Rα^-/-^ mice. **C.** Representative FCM analysis of liver mononuclear cells from NSG, NOD-SCID and NOD-SCID IL15Rα^-/-^ mice. **D.** Frequency of DX5^+^CD335^+^ murine NK cells among of CD3^-^B220^-^ cells from NSG (*n = 3 blood, n =3 liver, n = 2 spleen*) (magenta circles), NOD-SCID (*n = 5 blood, n = 5 liver, n = 6 spleen*) (orange squares), or NOD-SCID IL15Rα^-/-^ mice (*n = 5 blood, n = 6 liver, n = 7 spleen*) (green triangles). *** = p < 0.0005 by paired t-test. Mean ± SEM are shown. Data are from five single experiment.

### Results 1.2 NK cells from NOD-SCID IL15Rα^-/-^ mice have reduced tumor killing function

We performed a cytotoxicity assay to compare murine NK function from NOD-SCID IL15Rα^-/-^ mice (*n*=*3*) to NOD-SCID control mice (*n*=*3*). NK cells are the main cytotoxic population in NOD-SCID mice [32] and are functionally impaired in NSG mice due to *γ_c_* deficiency. However, NK cells were greatly reduced in NOD-SCID IL15Rα^-/-^ mice (Figure 1C and 1D) and we were unable to isolate sufficient NK cells from them for functional assays. Therefore, we co-cultured whole splenocyte or liver lymphocyte suspensions with CFSE-labelled non-target control cells from spleen or liver of NSG mice and e450-labelled target RLĴ1 murine lymphoma cells. Cytotoxicity of splenic cells, but not mononuclear cells from liver of NOD-SCID IL15Rα^-/-^ mice was significantly lower in comparison to that of NOD-SCID littermates (Figure 2A and 2B).

**Figure 2.**
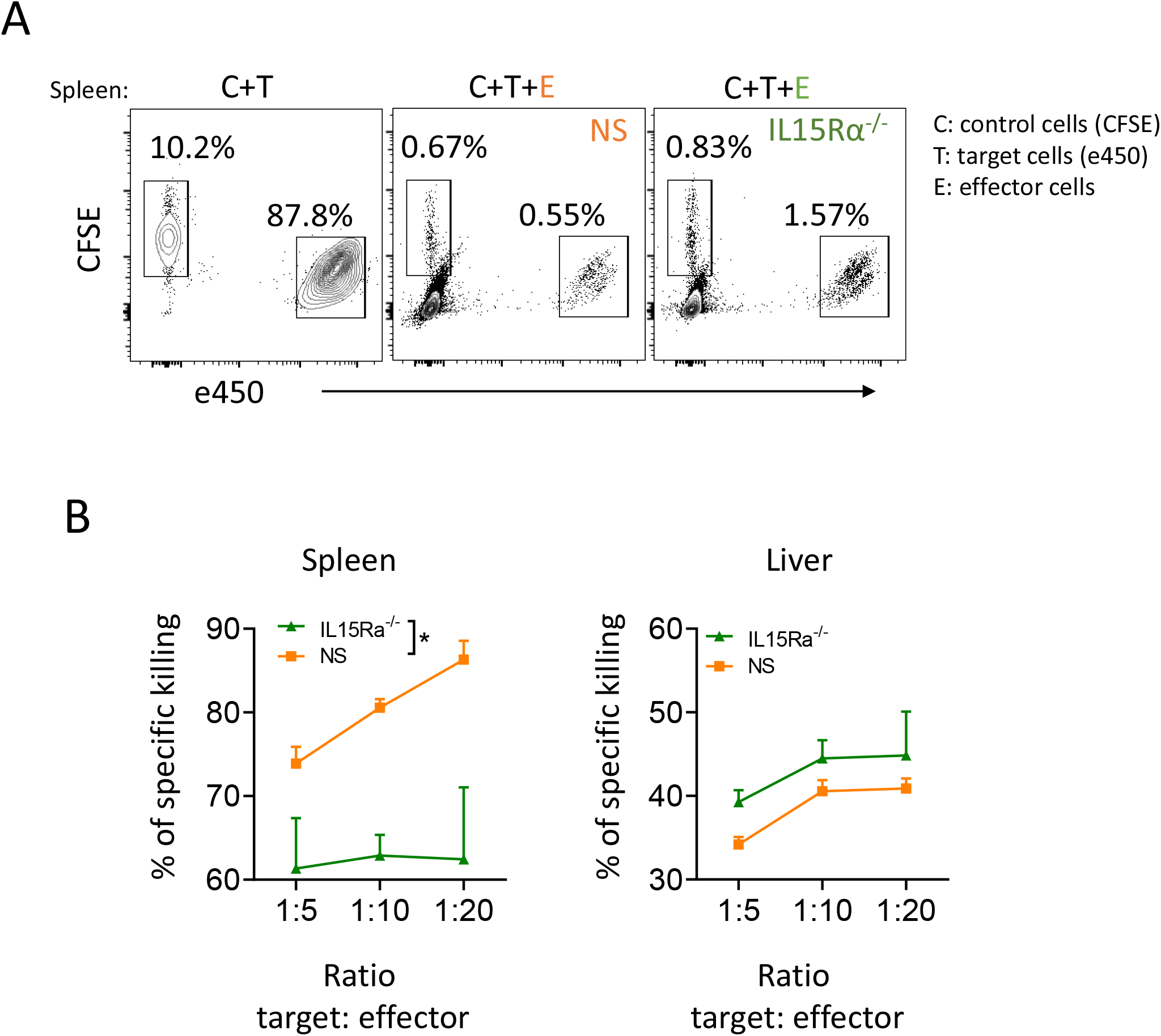
NK cells in NOD-SCID ILl5Rα^-/-^ mice show reduced cytotoxic function. NK cytotoxicity was assessed using a flow cytometry-based killing assay. Control cells (NSG splenocytes) and RL♂1 target cells were labeled with amine-binding dyes CFSE and e450. Cells from age and sex-matched NOD-SCID (labeled orange) (*n = 3*) or NOD-SCID IL15Rα^-/-^ (labeled green) mice (*n = 3*) were harvested and added with the control/target mixture at the indicated ratios. Splenocytes were incubated together for 4 hours at 37°C with 5% CO_2_, then stained with DAPI and analyzed. **A.** Representative FCM analysis and **B.** Percentage of specific killing calculated as Cytotoxicity = 100% × (1 - (Control/Target)_without effector_/(Control/Target)_with effector_). * = p < 0.05 by parametric paired t-test. Mean + SEM are shown. Data are from a single experiment.

### Results 1.3 Thymectomy does not prevent thymic lymphoma of NOD-SCID IL15Rα^-/-^ mice

NOD-SCID mice have a high incidence of thymic lymphoma that typically becomes evident around 20 weeks of age [8],[9]. By pooling from 5 independent experiments, we observed that NOD-SCID IL15Rα^-/-^ mice had shorter lifespans than did NOD-SCID mice (Figure 3A) and detected fulminant thymic lymphoma in peripheral blood, enlarged thymus and splenomegaly upon necropsy of both NOD-SCID and NOD-SCID IL15Rα^-/-^ strains of mice (Figure 3B). Many of these thymic cells expressed mouse CD4 and/or CD8 (Figure 3C and 3D) as well as CD3 (Supplementary Figure 2A and 2B) in NOD-SCID IL-15Rα^-/-^ mice. In an effort to reduce the incidence of thymic lymphoma, we surgically removed the mouse thymus from young (4-8-week-old) mice, but shorter survival was observed compared to thymectomized NOD-SCID littermates (Figure 3E).

**Figure 3.**
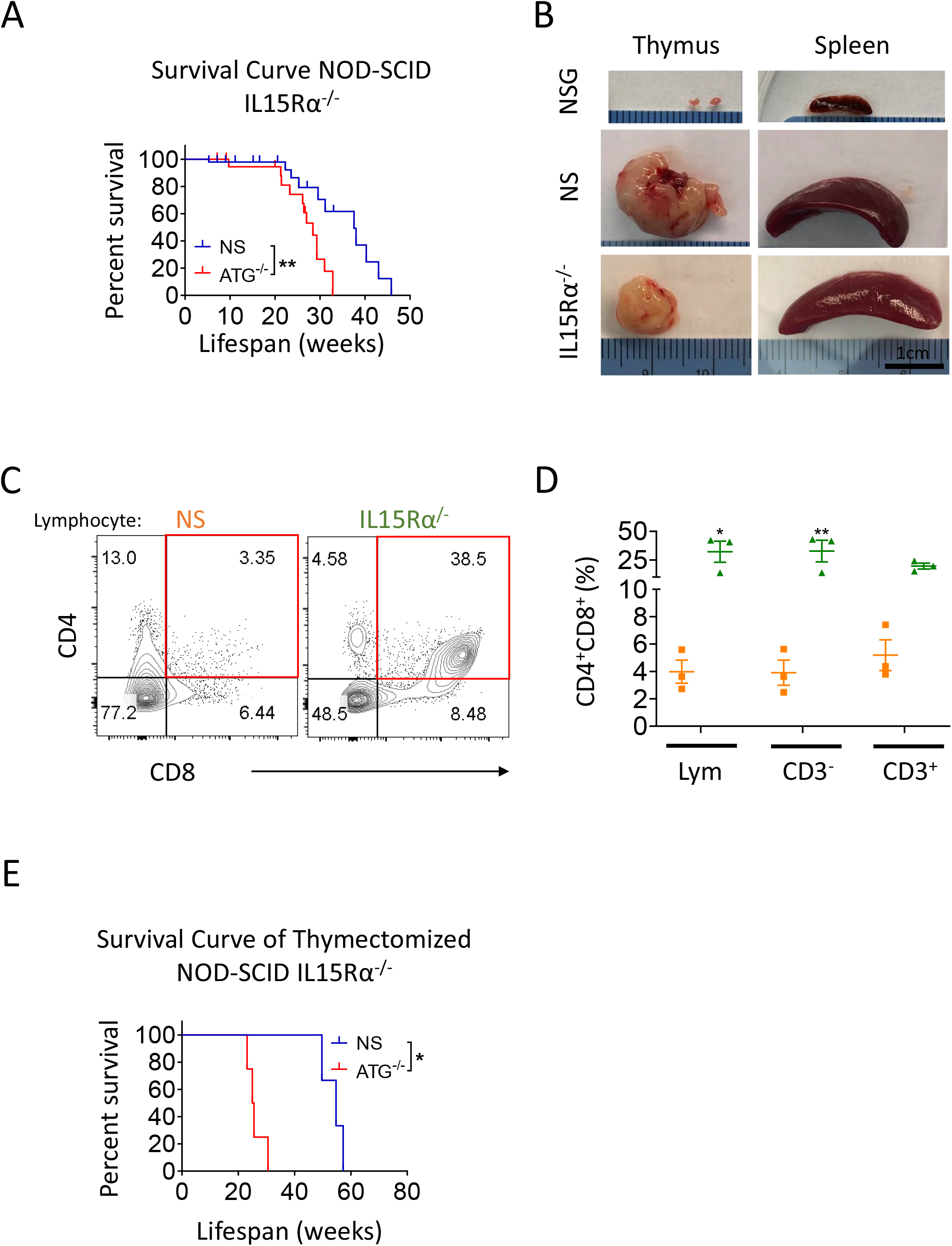
Lifespan of NOD-SCID IL15Rα^-/-^ mice is reduced and NOD-SCID IL15Rα^-/-^ mice develop fulminant thymic lymphomas. Kaplan-Meier survival curve comparing lifespan of NOD-SCID (*n = 48*) and NOD-SCID IL15Rα^-/-^ (*n = 25*) littermates. Data was pooled from 5 independent experiments. **B.** Representative images of thymic lymphoma tumors and enlarged spleens of 8 week-old NSG, NOD-SCID and NOD-SCID IL15Rα^-/-^ littermates. **C.** Representative FCM staining and **D.** Frequency of CD4^+^CD8^+^ thymic lymphoma cells in NOD-SCID (orange squares), or NOD-SCID IL15Rα^-/-^ mice (green triangles). **E.** Kaplan-Meier survival curves comparing the lifespan of NOD-SCID (*n = 3*) and NOD-SCID IL15Rα^-/-^ mice (*n = 4*) after surgical thymus removal at 4-6 weeks old. Data are from a single experiment. * = p < 0.05 by the Log-rank Mantel-Cox test for survival curve analysis. * = p < 0.05, ** = p < 0.005 by Bonferroni multiple comparison test. Mean ± SEM are shown. The term ‘’lymp” stands for lymphocyte population.

### Results 1.4 Human HSC engraftment of NOD-SCID IL15Rα^-/-^ mice

We used HIS NOD-SCID IL15Rα^-/-^, NOD-SCID and NSG mice to investigate if human immune cell engraftment improved due to the reduction of NK cells in NOD-SCID IL15Rα^-/-^ mice. We irradiated and transplanted 6-8-week old NSG, NOD-SCID, and NOD-SCID IL15Rα^-/-^ mice with human FL CD34^+^ cells and autologous human fetal thymus tissue. Surprisingly, NOD-SCID IL15Rα^-/-^ mice had the lowest level of human chimerism in peripheral blood, as well as in BM, LN and spleen compared to NSG and NOD-SCID littermates (Figure 4A and 4C). BM, LN and spleen were collected 18 weeks post-transplant from NSG, NOD-SCID and NOD-SCID IL15Rα^-/-^ recipients. No differences in total numbers of lymphocytes were found (Supplementary Figure 3A). However, human T cells and B cells were almost undetectable and essentially all human cells were monocytes in NOD-SCID IL15Rα^-/-^ mice (Figure 4D, 4E and 4F, and Supplementary Figure 3B, 3C and 3D), suggesting that resistance to human engraftment was somewhat selective for cells in the lymphoid lineages. Murine lymphoma cells were not detected in the blood of transplanted mice by FCM analysis (Supplementary Figure 3E). Altogether, these data show the absence of murine IL15Rα, did not improve human cell engraftment in NOD-SCID mice.

**Figure 4.**
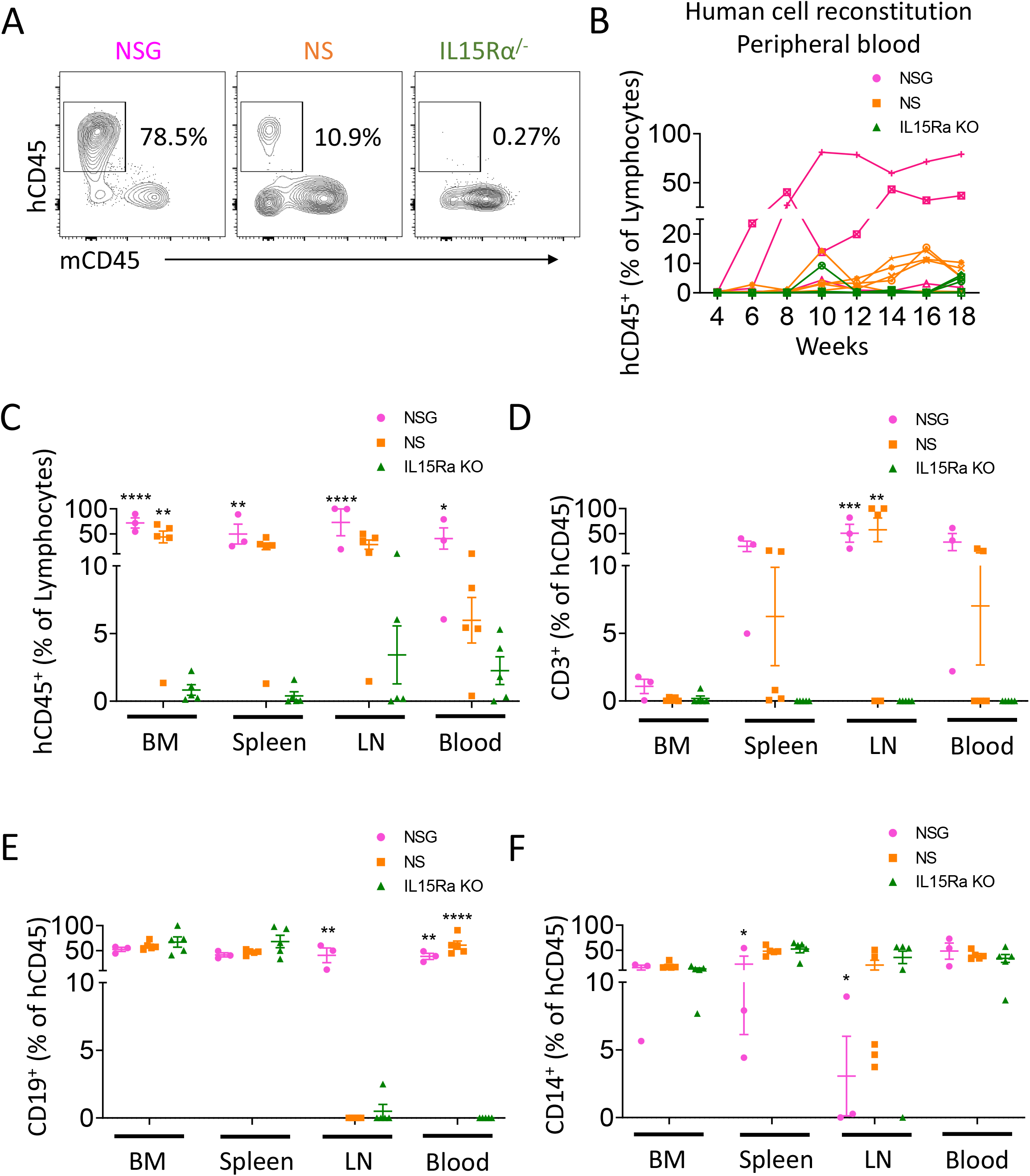
Poor engraftment of human cells in NOD-SCID IL15Rα^-/-^ mice. 6-8-week-old NSG (*n = 3*) and thymectomized NOD-SCID (*n = 5*) or NOD-SCID IL15Rα^-/-^ (*n = 5*) mice received 1 Gy total body irradiation and were transplanted intravenously with 2×10^5^ human fetal liver CD34^+^ cells and a fragment of cryopreserved autologous human fetal thymus was grafted under the renal capsule. **A.** Representative FCM staining of human and murine CD45^+^ cells in the white blood cells of transplanted mice 18 weeks post-transplantation. **B.** Human chimerism kinetics showing the percent of huCD45^+^ cells among WBCs in each mouse post-transplantation. **C.** Frequency of human CD45^+^ cells, **D.** CD3^+^ T cells, **E.** CD19^+^ and **F.** CD14^+^ monocytes in the bone marrow (BM), spleen, lymph nodes (LN) and peripheral blood 18 weeks post-humanization. * = p < 0.05, ** = p < 0.005, *** = p < 0.0005 by Bonferroni multiple comparison test. NSG (magenta circles), NOD-SCID (orange squares), or NOD-SCID IL15Rα^-/-^ mice (green triangles). Mean ± SEM are shown. Data are from a single experiment.

## Discussion

The initial identification of the SCID mutation in mice sparked the development of HIS mouse modeling [33],[34]. Decades of research have gone into developing precisely-engineered mice that are immune deficient while providing appropriate cytokine and physical support for engraftment of human hematopoietic stem cells and maintaining sufficient longevity to permit extended experimentation [35]–[37]. The goal of our study was to develop a more permissive mouse strain that retained lymphoid development to model the functions of human tissues and cells. We set out to develop an optimal host animal that combined the advantages of the NSG mouse strain with those of the NOD-SCID, which support reduced human peripheral blood chimerism but which have been shown to develop more robust secondary lymphoid structures than NSG mice [20]. The improved lymphoid structures in NOD-SCID mice are thought to occur because NOD-SCID animals have an intact γc receptor, which permits signaling of numerous γ_c_ cytokines such as IL-7. This cytokine signaling supports the development of lymphoid tissue inducer cells, critical stromal cells for secondary lymphoid development. However, murine NK cells in NOD-SCID animals reject transplanted human hematopoietic cells, leading to poor engraftment [12]. It is known that IL15 and IL15Rα signaling are required for mature NK cell development in mice [24],[38]. Therefore, it was reasonable to hypothesize that depleting NK cells on the NOD-SCID background by eliminating IL15Rα would generate an immunodeficient mouse strain that lacked NK cells and had improved lymphoid structure compared to NSG mice. We used CRISPR-Cas9 to delete the *IL15RA* gene. As expected, this deletion efficiently depleted murine NK cells on the NOD-SCID background. Interestingly, loss of cytotoxic function was detected in the spleen but not in the liver of NOD-SCID IL15Rα^-/-^ mice compared to NOD-SCID mice. This difference might be due to the heterogeneous nature of NK cells, as splenic NK cells predominantly express a mature phenotype, whereas liver NK cells express low levels of markers associated with maturation, including DX5, Mac-1 and Ly49 [39]. Maturation of immature NK cells might depend on the interaction between IL-15 and IL-15R in the spleen [40]–[42]. Therefore, the absence of IL-15Rα in NOD-SCID mice might have not affected the presence of immature NK cells in the liver. Immature liver NK cells have been shown to be quite cytotoxic [44–47] and in NOD-SCID IL15Rα^-/-^ mice could still mediate cytotoxicity via alternate killing mechanisms including IL-15 primed ligation of TNF-related apoptosis-inducing ligand (TRAIL) receptors [48],[49]. Although we removed IL-15Rα, IL-15 is still present in NOD-SCID IL15Rα^-/-^ and may bind to IL-2R to allow survival of liver immature NK cells. IL-15 uses components of the IL-2R, namely the β and γ chains, to bind with intermediate affinity and activate similar JAK/STAT pathways as IL-2 [50]–[51]. We believe that loss of IL15Rα in NOD-SCID mice might specifically affect the maturation and function of NK cells found in the spleen, and that the liver remains enriched with immature NK cells [52].

However, deleterious health effects were seen in NOD-SCID IL15Rα^-/-^ animals, which had significantly shorter lifespans than did NOD-SCID mice. NOD-SCID mice have long been known to develop a T cell thymic lymphoma [8],[9], and this tumor formation appears to be more rapid in the absence of murine NK cells, suggesting that NK cells play an important role in keeping these tumors at bay in NOD-SCID mice. We surgically removed the thymus from immunodeficient mice at early ages, hypothesizing that early surgical removal of thymus from NOD-SCID IL15Rα^-/-^ mice would reduce tumor development. The removal of murine thymus from NOD-SCID IL15Rα^-/-^ mice did appear to reduce thymic tumor burden, as we did not observe any local tumor formation. However, this removal did not prevent early mortality. The reduced lifespans of these mice may limit their utility as hosts for transplantation. Consistent with our observation of tumor development in NOD-SCID IL15Rα^-/-^ mice, NK cell depletion by absence of IL-15 cytokine or treatment with anti-asialo-GM1 mAb accelerated tumor development in B6 mice and reduced survival in NOD-SCID mice, respectively [53]. Since IL-15R β and γ chains and IL-15 are available in IL15Rα^-/-^ KO mice, our results suggest that high affinity IL-15Rα-dependent mature NK cells may play a role in protecting against these tumors in NOD-SCID mice.

We assessed whether the NOD-SCID IL15Rα^-/-^ mice could support improved engraftment of transplanted human CD34^+^ cells compared to NOD-SCID or NSG mice. Unexpectedly, our results showed that human engraftment in NOD-SCID IL15Rα^-/-^ mice was negligible, lower even than that seen in NOD-SCID mice. This result was not predictable based on anything that has been reported about the IL15/IL15Rα signaling pathway and points to novel biological functions for IL15Rα that have not been previously explored. IL15 and IL15Rα have primarily been studied in the context of CD8 memory and NK cell development [24], although studies have implicated it in muscle metabolism [25] and bone formation [54],[55]. The results shown here, indicate that IL15Rα-deficient animals are nearly incapable of supporting human T and B cell engraftment. Since cytotoxic activity in the liver was similar in IL15Rα^-/-^, and NOD-SCID mice, we hypothesize that immature NK cells that can receive IL-15 signals through the intermediate affinity receptor may be capable of rejecting human hematopoietic cells.

Ultimately, our results have shown that the NOD-SCID IL15Rα^-/-^ mouse strain has poor viability and supports negligible human engraftment and is thus unlikely to be optimal for future human transplantation studies. However, our results also suggest that further studies using these animals may be of value. Our results reinforce the importance of NK cells in anti-tumor immunity and provide a model system for investigating mechanisms of human hematopoietic cell rejection in greater detail.

## Acknowledgements

This work was supported by National Institutes of Health (NIH) grants #P01AI045897 and NIDDK grants #UC4DK104207, #U01DK123559, #R01DK103585 as well as the T1D Resource grant # DK097610. MVW and BEL were also supported in part by The Jackson Laboratory. These studies used the resources of the Diabetes Research Center Flow Core Facility funded in part through Center Grant #P30DK063608, including LSRII instrument acquired with support from award #S10RR027050. We gratefully acknowledge the contribution of Microinjection Services at The Jackson Laboratory for expert assistance with the work described in this publication.

## Authorship

Contribution: AV and GN designed and performed experiments, analyzed results and wrote the manuscript; MKM, ND, HL and EC participated in the performance of experiments. MCK performed removal of thymus in mice. ND assisted in writing the manuscript. EC assisted with animal husbandry and genotyping. BEL, MVW and DS performed gene editing, generated the animal model for this study, provided intellectual input and assisted write the paper. LD designed sgRNAs and provided advice on model development. MS oversaw the design, conduct and interpretation of all experiments, and with AV and GN, wrote the manuscript.

The authors declare that they have no conflict of interests.

---

**Supplementary Figure 1.**
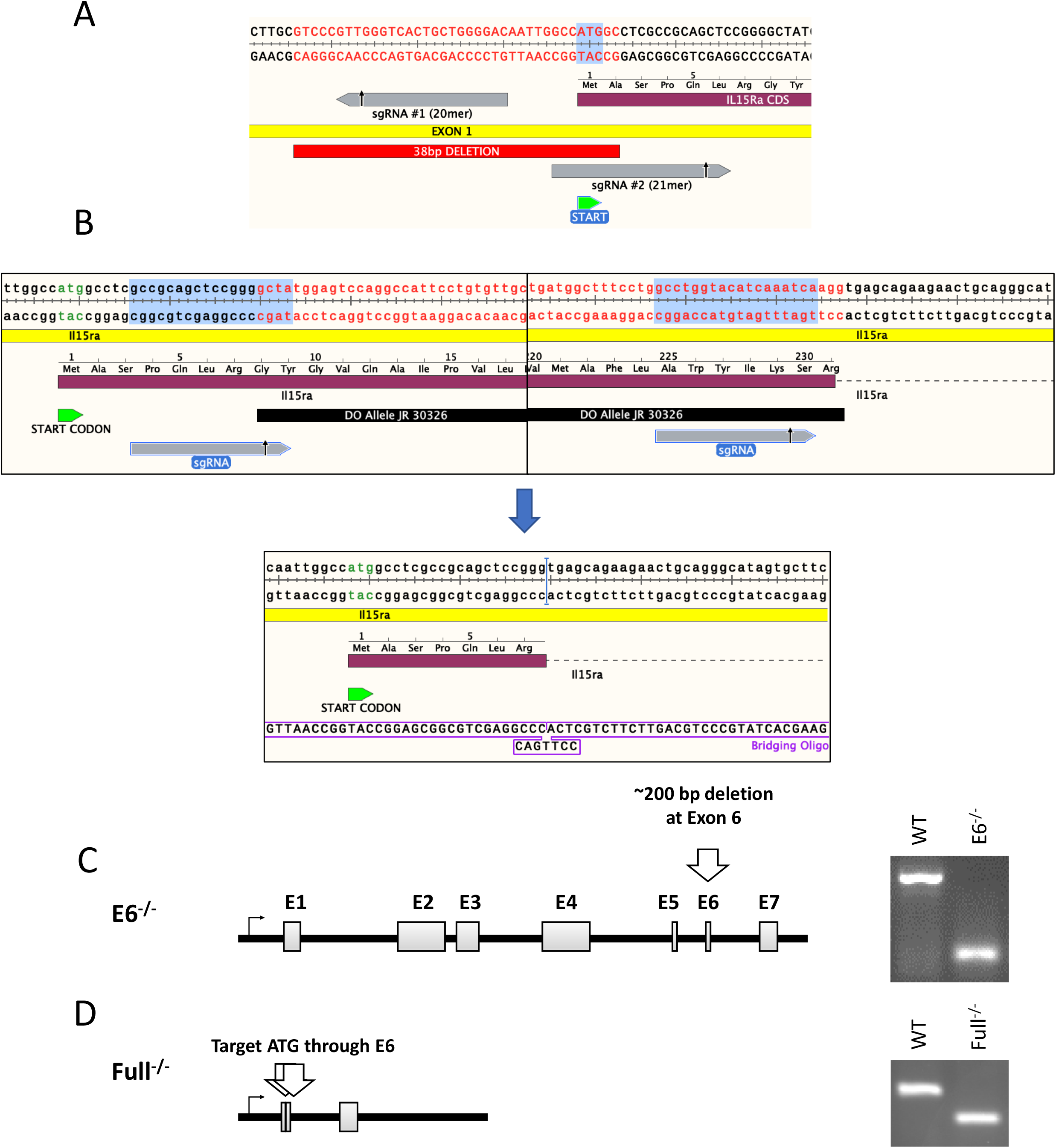
CRISPR/CAS9 targeting of IL15RA locus by targeting ATG start codon. **A.** Diagram of the *IL15RA* locus and 38bp depletion in exon 1 using sgRNA #1 (20mef) and sgRNA #2 (21mer). CRISPR target sites near the start codon (ATG) and 38bp deletion. **B.** Diagram of the *IL15RA* locus and 24,778bp depletion resulting from dual sgRNA. **C.** CRISPR target sites near *IL15RA* exon 6 **D.** CRISPR target sites *IL15RA* start codon (ATG) and exon 6 together.

**Supplementary Figure 2.**
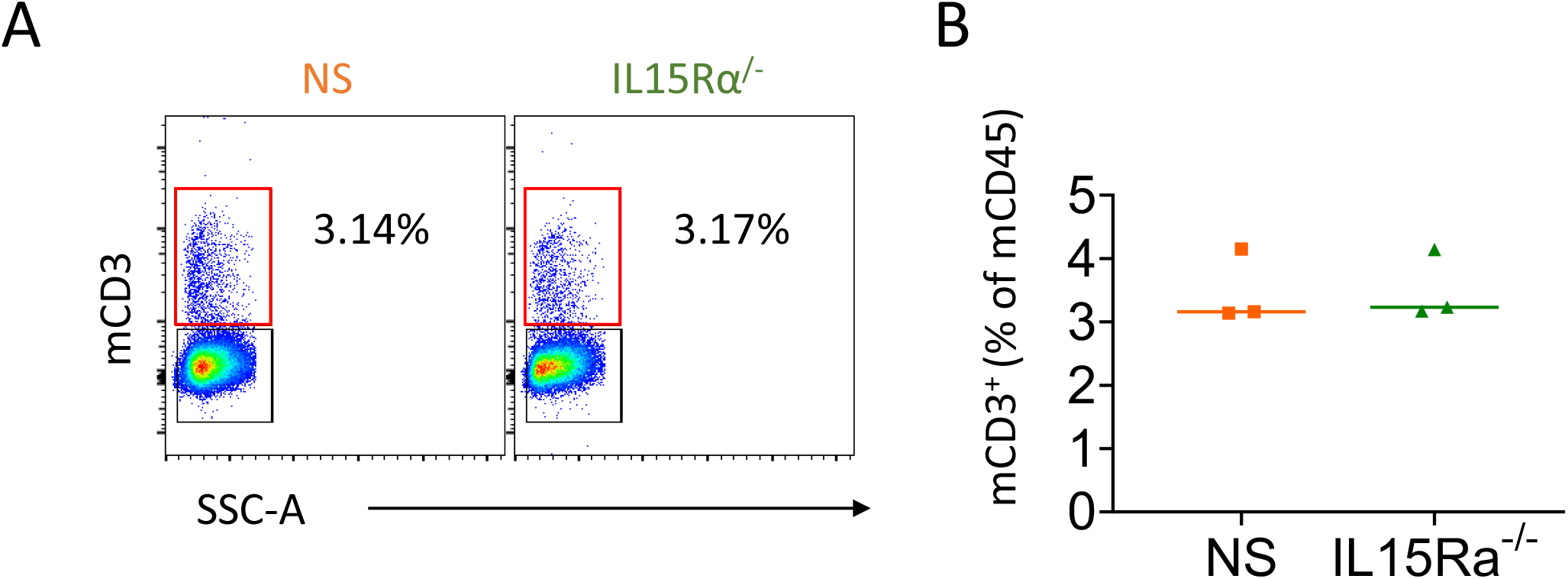
Detection of mouse T cells in native thymus of NOD-SCID IL15Rα^-/-^ mice. **A.** Representative FCM staining and **B.** Frequency of mouse CD3^+^ T cells. NOD-SCID (*n = 3*) (orange squares), or NOD-SCID IL15Rα^-/-^ mice (*n = 3*) (green triangles). Mean is shown. Data are from a single experiment.

**Supplementary Figure 3.**
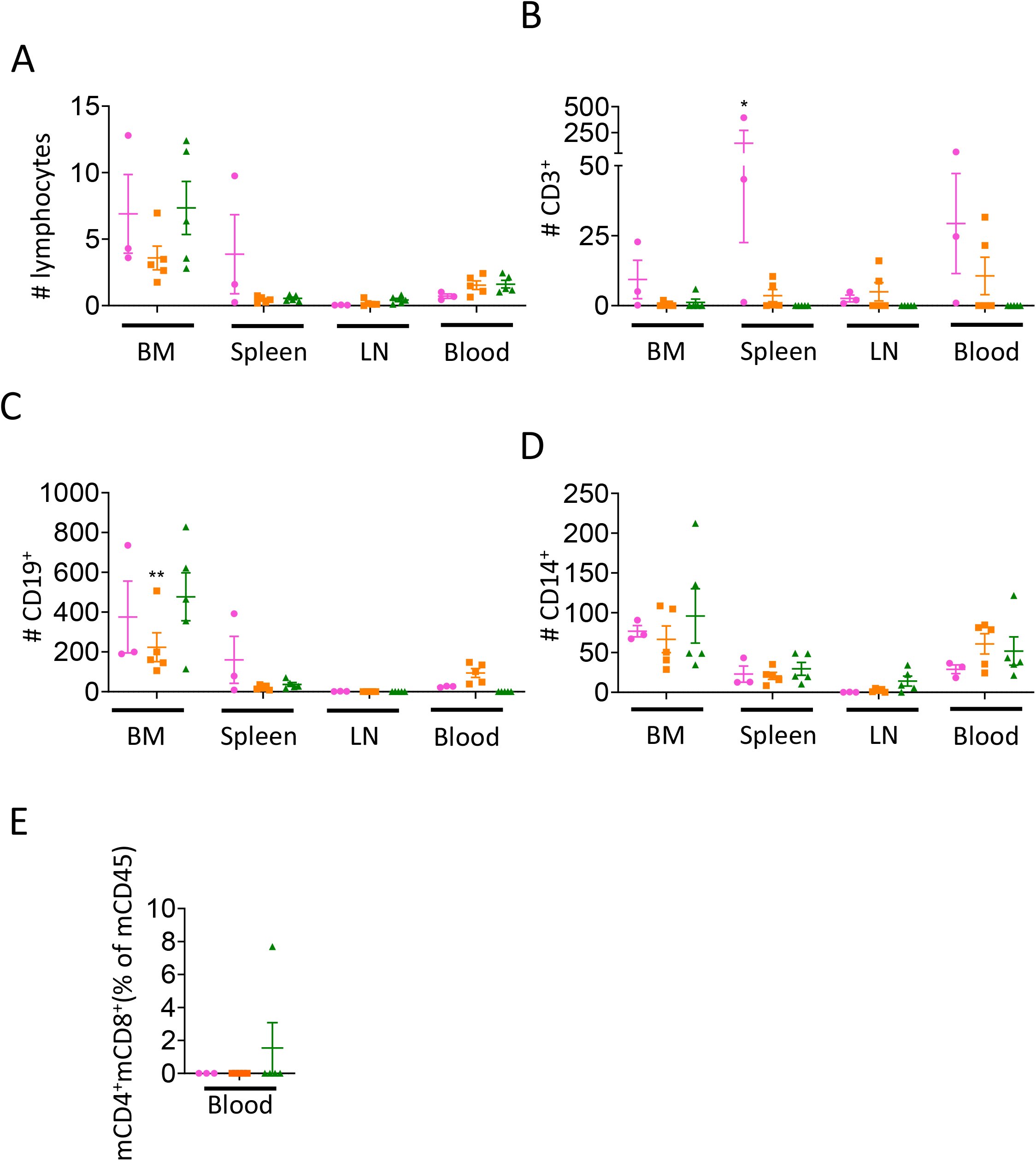
Human T cell, B cell and monocyte chimerism. **A.** Total number of **B.** CD3^+^ T cells, **C.** CD19^+^ B cells and **D.** CD14^+^ monocytes within human CD45^+^ cells in bone marrow (one leg), spleen, lymph nodes and peripheral blood (200uL). * = p < 0.05, ** = p < 0.005, *** = p < 0.0005 by Bonferroni multiple comparison test. **D.** Murine thymic lymphoma cells are not detected in blood of transplanted mice by FCM. NSG (*n = 3*) (magenta circles), NOD-SCID (*n = 5*) (orange squares), or NOD-SCID IL15Rα^-/-^ mice (*n = 5*) (green triangles). Mean ± SEM are shown. Data are from a single experiment.

## Reference

1. Lan P, Tonomura N, Shimizu A, Wang S, Yang YG. Reconstitution of a functional human immune system in immunodeficient mice through combined human fetal thymus/liver and CD34+ cell transplantation. Blood. 2006.DOI: 10.1182/blood-2005-11-4388.

2. Aryee KE, Shultz LD, Brehm MA. Immunodeficient mouse model for human hematopoietic stem cell engraftment and immune system development. Methods Mol. Biol. 2014.DOI: 10.1007/978-1-4939-1133-2_18.

3. Greiner DL, Hesselton RA, Shultz LD. SCID mouse models of human stem cell engraftment. Stem Cells. 1998.DOI: 10.1002/stem.160166.

4. Ito R, Takahashi T, Katano I, Ito M. Current advances in humanized mouse models. Cell. Mol. Immunol. 2012; 9:208–14. Available at: http://www.pubmedcentral.nih.gov/articlerender.fcgi?artid=4012844&tool=pmcentrez&rendertype=abstract.DOI: 10.1038/cmi.2012.2.

5. Kwong LS, Brown MH, Barclay AN, Hatherley D. Signal-regulatory protein α from the NOD mouse binds human CD47 with an exceptionally high affinity - implications for engraftment of human cells. Immunology. 2014.DOI: 10.1111/imm.12290.

6. Legrand N, Huntington ND, Nagasawa M, Bakker AQ, Schotte R, Strick-Marchand H, De Geus SJ, et al. Functional CD47/signal regulatory protein alpha (SIRPα) interaction is required for optimal human T- and natural killer-(NK) cell homeostasis in vivo. Proc. Natl. Acad. Sci. U. S. A. 2011.DOI: 10.1073/pnas.1101398108.

7. Wong ASL, Mortin-Toth S, Sung M, Canty AJ, Gulban O, Greaves DR, Danska JS. Polymorphism in the Innate Immune Receptor SIRPα Controls CD47 Binding and Autoimmunity in the Nonobese Diabetic Mouse. J. Immunol. 2014.DOI: 10.4049/jimmunol.1401984.

8. Serreze D V., Leiter EH, Hanson MS, Christianson SW, Shultz LD, Hesselton RM, Greiner DL. Emv30(null) NOD-scid mice: An improved host for adoptive transfer of autoimmune diabetes and growth of human lymphohematopoietic cells. Diabetes. 1995; 44:1392–1398. DOI: 10.2337/diab.44.12.1392.

9. Prochazka M, Gaskins HR, Shultz LD, Leiter EH. The nonobese diabetic scid mouse: Model for spontaneous thymomagenesis associated with immunodeficiency. Proc. Natl. Acad. Sci. U. S. A. 1992.DOI: 10.1073/pnas.89.8.3290.

10. Shultz LD, Lyons BL, Burzenski LM, Gott B, Chen X, Chaleff S, Kotb M, et al. Human lymphoid and myeloid cell development in NOD/LtSz-scid IL2R gamma null mice engrafted with mobilized human hemopoietic stem cells. J. Immunol. 2005; 174:6477–6489. DOI: 10.4049/jimmunol.174.10.6477.

11. Denton PW, Nochi T, Lim A, Krisko JF, Martinez-Torres F, Choudhary SK, Wahl A, et al. IL-2 receptor γ-chain molecule is critical for intestinal T-cell reconstitution in humanized mice. Mucosal Immunol. 2012.DOI: 10.1038/mi.2012.31.

12. Ito M, Hiramatsu H, Kobayashi K, Suzue K, Kawahata M, Hioki K, Ueyama Y, et al. NOD/SCID/γ cnull mouse: An excellent recipient mouse model for engraftment of human cells. Blood. 2002.DOI: 10.1182/blood-2001-12-0207.

13. Russell SM, Keegan AD, Harada N, Nakamura Y, Noguchi M, Leland P, Friedmann MC, et al. Interleukin-2 receptor γ chain: A functional component of the interleukin-4 receptor. Science (80-.). 1993. DOI: 10.1126/science.8266078.

14. Noguchi M, Nakamura Y, Russell SM, Ziegler SF, Tsang M, Cao X, Leonard WJ. Interleukin-2 receptor γ chain: A functional component of the interleukin-7 receptor. Science (80-.). 1993.DOI: 10.1126/science.8266077.

15. Asao H, Okuyama C, Kumaki S, Ishii N, Tsuchiya S, Foster D, Sugamura K. Cutting Edge: The Common γ-Chain Is an Indispensable Subunit of the IL-21 Receptor Complex. J. Immunol. 2001.DOI: 10.4049/jimmunol.167.1.1.

16. Rochman Y, Spolski R, Leonard WJ. New insights into the regulation of T cells by γc family cytokines. Nat. Rev. Immunol. 2009. DOI: 10.1038/nri2580.

17. Cairns R, Papandreou I, Denko N. Overcoming physiologic barriers to cancer treatment by molecularly targeting the tumor microenvironment. Mol. Cancer Res. 2006; 4:61–70. Available at: http://www.ncbi.nlm.nih.gov/pubmed/16513837.DOI: 10.1158/1541-7786.MCR-06-0002.

18. Brehm MA, Shultz LD, Luban J, Greiner DL. Overcoming Current Limitations in Humanized Mouse Research. J. Infect. Dis. 2013.DOI: 10.1093/infdis/jit319.

19. Chappaz S, Finke D. The IL-7 Signaling Pathway Regulates Lymph Node Development Independent of Peripheral Lymphocytes. J. Immunol. 2010.DOI: 10.4049/jimmunol.0901647.

20. Nochi T, Denton PW, Wahl A, Garcia JV. Cryptopatches Are Essential for the Development of Human GALT. Cell Rep. 2013.DOI: 10.1016/j.celrep.2013.05.037.

21. Yoshino H, Ueda T, Kawahata M, Kobayashi K, Ebihara Y, Manabe A, Tanaka R, et al. Natural killer cell depletion by anti-asialo GM1 antiserum treatment enhances human hematopoietic stem cell engraftment in NOD/Shi-scid mice. Bone Marrow Transplant. 2000.DOI: 10.1038/sj.bmt.1702702.

22. Schluns KS, Stoklasek T, Lefrançois L. The roles of interleukin-15 receptor α: Trans-presentation, receptor component, or both? Int. J. Biochem. Cell Biol. 2005.DOI: 10.1016/j.biocel.2005.02.017.

23. Ali AK, Nandagopal N, Lee SH. IL-15-PI3K-AKT-mTOR: A critical pathway in the life journey of Natural Killer cells. Front. Immunol. 2015.DOI: 10.3389/fimmu.2015.00355.

24. Kennedy MK, Glaccum M, Brown SN, Butz EA, Viney JL, Embers M, Matsuki N, et al. Reversible defects in natural killer and memory CD8 T cell lineages in interleukin 15-deficient mice. J. Exp. Med. 2000.DOI: 10.1084/jem.191.5.771.

25. Pistilli EE, Bogdanovich S, Garton F, Yang N, Gulbin JP, Conner JD, Anderson BG, et al. Loss of IL-15 receptor α alters the endurance, fatigability, and metabolic characteristics of mouse fast skeletal muscles. J. Clin. Invest. 2011.DOI: 10.1172/JCI44945.

26. Loro E, Ramaswamy G, Chandra A, Tseng WJ, Mishra MK, Shore EM, Khurana TS. IL15RA is required for osteoblast function and bone mineralization. Bone. 2017.DOI: 10.1016/j.bone.2017.06.003.

27. Cong L, Ran FA, Cox D, Lin S, Barretto R, Habib N, Hsu PD, et al. Multiplex genome engineering using CRISPR/Cas systems. Science. 2013; 339:819–823. DOI: 10.1126/science.1231143.

28. Low BE, Kutny PM, Wiles M V. Simple, efficient CRISPR-cas9-mediated gene editing in mice: Strategies and methods. Methods Mol. Biol. 2016.DOI: 10.1007/978-1-4939-3661-8_2.

29. Borsotti C, Danzl NM, Nauman G, Hölzl MA, French C, Chavez E, Khosravi-Maharlooei M, et al. HSC extrinsic sex-related and intrinsic autoimmune disease–related human B-cell variation is recapitulated in humanized mice. Blood Adv. 2017.DOI: 10.1182/bloodadvances.2017006932.

30. Khosravi-Maharlooei M, Hoelzl M, Li HW, Madley RC, Waffarn EE, Danzl NM, Sykes M. Rapid thymectomy of NSG mice to analyze the role of native and grafted thymi in humanized mice. Eur. J. Immunol. 2020.DOI: 10.1002/eji.201948205.

31. Li HW, Vishwasrao P, Hölzl MA, Chen S, Choi G, Zhao G, Sykes M. Impact of Mixed Xenogeneic Porcine Hematopoietic Chimerism on Human NK Cell Recognition in a Humanized Mouse Model. Am. J. Transplant. 2017.DOI: 10.1111/ajt.13957.

32. Christianson SW, Greiner DL, Hesselton RA, Leif JH, Wagar EJ, Schweitzer IB, Rajan T V, et al. Enhanced human CD4+ T cell engraftment in beta2-microglobulin-deficient NOD-scid mice. J. Immunol. 1997.

33. McCune JM, Namikawa R, Kaneshima H, Shultz LD, Lieberman M, Weissman IL. The SCID-hu mouse: murine model for the analysis of human hematolymphoid differentiation and function. Science (80-.). 1988; 241:1632–9. Available at: http://www.ncbi.nlm.nih.gov/pubmed/2971269.DOI: 10.1126/science.2971269.

34. Hayes SM, Greiner DL, Shultz LD, Greiner DL. Thymic Involution in Viable Motheaten (mev) Mice is Associated with a Loss of Intrathymic Precursor Activity. Dev. Immunol. 1992; 2:191–205. DOI: 10.1155/1992/68954.

35. Willinger T, Rongvaux A, Strowig T, Manz MG, Flavell RA. Improving human hemato-lymphoid-system mice by cytokine knock-in gene replacement. Trends Immunol. 2011; 32:321–327. DOI: 10.1016/j.it.2011.04.005.

36. Billerbeck E, Barry WT, Mu K, Dorner M, Rice CM, Ploss A. Development of human CD4+FoxP3+ regulatory T cells in human stem cell factor-, granulocyte-macrophage colony-stimulating factor-, and interleukin-3-expressing NOD-SCID IL2Rγ(null) humanized mice. Blood. 2011; 117:3076–3086. DOI: 10.1182/blood-2010-08-301507.

37. Yoshihara S, Li Y, Xia J, Danzl N, Sykes M, Yang YG. Posttransplant hemophagocytic lymphohistiocytosis driven by myeloid cytokines and vicious cycles of T-cell and macrophage activation in humanized mice. Front. Immunol. 2019; 10:1–10. DOI: 10.3389/fimmu.2019.00186.

38. Meazza R, Azzarone B, Orengo AM, Ferrini S. Role of Common-Gamma Chain Cytokines in NK Cell Development and Function: Perspectives for Immunotherapy. 2011; 2011. DOI: 10.1155/2011/861920.

39. Kim S, Iizuka K, Kang HSP, Dokun A, French AR, Greco S, Yokoyama WM. In vivo developmental stages in murine natural killer cell maturation. Nat. Immunol. 2002. DOI: 10.1038/ni796.

40. Sojka DK, Plougastel-Douglas B, Yang L, Pak-Wittel MA, Artyomov MN, Ivanova Y, Zhong C, et al. Tissue-resident natural killer (NK) cells are cell lineages distinct from thymic and conventional splenic NK cells. Elife. 2014.DOI: 10.7554/eLife.01659.

41. Crinier A, Milpied P, Escalière B, Piperoglou C, Galluso J, Balsamo A, Spinelli L, et al. High-Dimensional Single-Cell Analysis Identifies Organ-Specific Signatures and Conserved NK Cell Subsets in Humans and Mice. Immunity. 2018.DOI: 10.1016/j.immuni.2018.09.009.

42. Kawamura T, Koka R, Ma A, Kumar V. Differential Roles for IL-15R α-Chain in NK Cell Development and Ly-49 Induction. J. Immunol. 2003.DOI: 10.4049/jimmunol.171.10.5085.

43. Takeda K, Cretney E, Hayakawa Y, Ota T, Akiba H, Ogasawara K, Yagita H, et al. TRAIL identifies immature natural killer cells in newborn mice and adult mouse liver. Blood. 2005.DOI: 10.1182/blood-2004-08-3262.

44. Tang L, Peng H, Zhou J, Chen Y, Wei H, Sun R, Yokoyama WM, et al. Differential phenotypic and functional properties of liver-resident NK cells and mucosal ILC1s. J. Autoimmun. 2016; 67:29–35. Available at: http://dx.doi.org/10.1016/j.jaut.2015.09.004.DOI: 10.1016/j.jaut.2015.09.004.

45. Zhou J, Peng H, Li K, Qu K, Wang B, Wu Y, Ye L, et al. Liver-Resident NK Cells Control Antiviral Activity of Hepatic T Cells via the PD-1-PD-L1 Axis. Immunity. 2019; 50:403–417.e4. Available at: https://doi.org/10.1016/j.immuni.2018.12.024.DOI: 10.1016/j.immuni.2018.12.024.

46. Nabekura T, Riggan L, Hildreth AD, O’Sullivan TE, Shibuya A. Type 1 Innate Lymphoid Cells Protect Mice from Acute Liver Injury via Interferon-γ Secretion for Upregulating Bcl-xL Expression in Hepatocytes. Immunity. 2020; 52:96–108.e9. Available at: https://doi.org/10.1016/j.immuni.2019.11.004.DOI: 10.1016/j.immuni.2019.11.004.

47. Chen Y, Tian Z. Innate lymphocytes: pathogenesis and therapeutic targets of liver diseases and cancer. Cell. Mol. Immunol. 2021; 18:57–72. Available at: http://dx.doi.org/10.1038/s41423-020-00561-z.DOI: 10.1038/s41423-020-00561-z.

48. Wagner JA, Rosario M, Romee R, Berrien-Elliott MM, Schneider SE, Leong JW, Sullivan RP, et al. CD56bright NK cells exhibit potent antitumor responses following IL-15 priming. J. Clin. Invest. 2017.DOI: 10.1172/JCI90387.

49. Ma A, Koka R, Burkett P. DIVERSE FUNCTIONS OF IL-2, IL-15, AND IL-7 IN LYMPHOID HOMEOSTASIS. Annu. Rev. Immunol. 2006.DOI: 10.1146/annurev.immunol.24.021605.090727.

50. Vámosi G, Bodnár A, Vereb G, Jenei A, Goldman CK, Langowski J, Tóth K, et al. IL-2 and IL-15 receptor α-subunits are coexpressed in a supramolecular receptor cluster in lipid rafts of T cells. Proc. Natl. Acad. Sci. U. S. A. 2004.DOI: 10.1073/pnas.0403916101.

51. Lin JX, Migone TS, Tseng M, Friedmann M, Weatherbee JA, Zhou L, Yamauchi A, et al. The role of shared receptor motifs and common stat proteins in the generation of cytokine pleiotropy and redundancy by IL-2, IL-4, IL-7, IL-13, and IL-15. Immunity. 1995.DOI: 10.1016/1074-7613(95)90141-8.

52. Peng H, Jiang X, Chen Y, Sojka DK, Wei H, Gao X, Sun R, et al. Liver-resident NK cells confer adaptive immunity in skin-contact inflammation. J. Clin. Invest. 2013.DOI: 10.1172/JCI66381.

53. Rauch DA, Harding JC, Ratner L. IL-15 deficient tax mice reveal a role for IL-1α in tumor immunity. PLoS One. 2014.DOI: 10.1371/journal.pone.0085028.

54. Ogata Y, Kukita A, Kukita T, Komine M, Miyahara A, Miyazaki S, Kohashi O. A Novel Role of IL-15 in the Development of Osteoclasts: Inability to Replace Its Activity with IL-2. J. Immunol. 1999; 162:2754 LP – 2760.

55. Takeda H, Kikuchi T, Soboku K, Okabe I, Mizutani H, Mitani A, Ishihara Y, et al. Effect of IL-15 and natural killer cells on osteoclasts and osteoblasts in a mouse coculture. Inflammation. 2014; 37:657–669. DOI: 10.1007/s10753-013-9782-0.

